# Auto-regressive Rank Order Similarity (aros) test

**DOI:** 10.1101/2022.06.15.496113

**Authors:** Tommy Clausner, Stefano Gentili

**Affiliations:** Lyon Neuroscience Research Center, Computation, Cognition and Neurophysiology (Cophy) team, INSERM UMRS 1028, CNRS UMR 5292, Université Claude Bernard Lyon 1, France

**Keywords:** rank order, multiple comparison, permutation test

## Abstract

In the present paper we propose a non-parametric statistical test procedure for interval scaled, paired samples data that circumvents the multiple comparison problem (MCP) by relating the data to the rank order of its group averages. Using an auto-regressive procedure, a single test statistic for multiple groups is obtained that allows for qualitative statements about whether multiple group averages are in fact different and how they can be sorted. The presented procedure outperforms classical tests, such as pairwise conducted t-tests and ANOVA, in some circumstances. Furthermore, the test is robust against noise and does not require the data to follow any particular distribution. If *A* is a data matrix containing *N* observations for *k* groups, then the test statistic *η* can be computed by 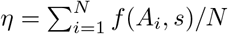, where **s** is a vector of length *k* containing the average for each group, transformed into unique rank values. This statistic is compared to the distribution *D*, obtained by Monte Carlo sampling from the permutation distribution. It will be demonstrated that *D* can be described by a normal distribution for a variety of input data distributions and choices for *f*, as long as a set of criteria is met. Comparing *η* to the permutation distribution controls the false alarm (FA) rate sufficiently, since the exact p-value can be estimated [1]. Multiple examples of possible choices for *f* will be discussed, as well as detailed descriptions of the underlying test assumptions, possible interpretations and use cases. All mathematical derivations are supported with a set of simulations, written in Python that can be downloaded from https://gitlab.com/TommyClausner/aros-test together with an implementation of the test itself.

## 1 Introduction

In this paper we propose a paired samples, nonparametric statistical procedure on the basis of a permutation test, which aims to circumvent the multiple comparison problem (MCP) by combining multiple group averages into a single statistical value. The MCP arises from the fact that in order to control the false alarm (FA) rate for comparing more than two groups, the critical *α* level needs to be adjusted. Commonly accepted levels for FA rates are less than 5% or 1%. This means that the null hypothesis (*H*_0_) was falsely rejected in less than 5% or 1% of all cases. As the number of groups *k* increase, the number of tests *n_t_* increases with *n_t_* = *k*!/(2(*k* – 2)!). If no adjustments to the critical *α* was made, the FA rate increases from ≤ 0.05 to ≤ 1 – (1 – *α*)^*n_t_*^, which for three groups means the FA rate becomes ≤ 0.14. This issue is commonly referred to as multiple comparison problem. One common strategy is to adjust the α level (lower it), until a satisfactory control for the FA rate is achieved ^1^. Adjusting the α level however comes at the cost of potentially reducing statistical power, that is the sensitivity of the test, and thus the rate at which the so called type II errors (falsely accepting *H*_0_) are produced increases.

A widely known test used for statistical analyses of more than two groups is the analysis of variances (ANOVA). The one-way ANOVA, for instance, uses an *F*-test to relate the variances within and between groups to each other. However, a significant result (i.e. rejecting the *H*_0_, which states no difference between the group means) only indicates a difference between any subset of the means. Thus, a significant result is often followed by a pairwise comparison, which again leads to the aforementioned MCP.

Furthermore, classical tests such as *z*-tests, t-tests or *F*-tests often require the underlying data to be normal distributed or at least that the parameters of the underlying distributions are known. Instead of relying on knowledge of the underlying distribution, the distribution under *H*_0_ can be estimated by a permutation procedure, which can be applied to all of the above classical tests. This procedure is well established for a variety of use cases [2–4] and allows for an exact estimation of the *p*-value [1]. The fundamental idea behind permutation tests is that if there is no difference between multiple averages, then the members of each group can be exchanged, because they stem from the same distribution (*H*_0_). Commonly, members of each group are shuffled between the groups and the respective test statistic of interest is computed. By repeating the procedure a large number of times (ideally by iterating through all possible combinations), the permutation distribution under *H*_0_ is obtained [5]. In a last step the test statistic of the original data is compared to the permutation distribution and the *p*-value is computed as the fraction of all values of the permutation distribution that exceed the test statistic computed from the original data [1]. Since the *p*-value is computed by Monte Carlo sampling from the permutation distribution, no knowledge about the underlying parameters is required [6]. Thus, permutation tests belong to the family of non-parametric tests.

The proposed auto-regressive rank order similarity (aros) test is constructed as a permutation test and thus belongs to the family of non-parametric tests as well. Additionally, the MCP is circumvented by condensing the relationship between the data and each group average into a single statistic. Precisely, the group data for each paired observation is treated as vector that is related to the vector of the rank order of the average group value, by means of a similarity metric. Thus, the test statistic *η* can be seen as the average similarity between the observations and the rank order of the averages. It is tested whether this similarity is significantly greater than the average similarity within the permutation distribution under *H*_0_.

## 2 The aros test explained

The aros test is a non-parametric statistical hypothesis test for paired sampled, interval scaled data for multiple group average comparison. However, as compared to conventional tests on interval scaled data, the aros test does not rely on the difference between the means of multiple distributions, but rather on the relationship of each paired observation within the data to an ordinal scaled profile or shape derived from either the group averages or an external source. If we assume *k* = 3 groups with averages *μ_a_, μ_b_, μ_c_*, the standard procedure for classical tests would be to pairwise compare those three groups in order to obtain the qualitative and quantitative relationship between those averages. However, when applying the aros test, this question is reduced to the qualitative relationship of how the group averages rank relative to each other and how well the data explains this relationship. This means that if we observe a ranking of the the means such that *μ_b_* < *μ_c_* < *μ_a_*, auto-regressive rank order similarity tests, whether this relationship can truly be justified by the data (as compared to determining, whether the pairwise difference between those means is zero in a more traditional setting). In other words it is tested, whether - on average - each observation expresses a similar relationship between each group or condition as expressed by the group averages. The most general formulation for the test statistic *η* can be written as:

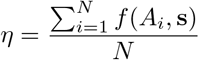

where **s** is the vector of group averages, transformed into a set of unique, evenly spaced rank values, such that **s** = [*s*_1_, …, *s_k_*]. For the aforementioned relationship *μ_b_* < *μ_c_* < *μ_a_*, one possibility would be to set **s** = [3, 1, 2]. The function *f* relates each observation *A*_*i*∈[1, …, *N*]_ (the values for each observation in all conditions) of length *k* and **s** separately. *η* is obtained by averaging the results from *f* (*A_i_*, **s**). Thereby, the function *f* can be freely chosen, as long as *f* relates *A_i_* and **s** in terms of similarity. Examples for *f* might be the least square solution to **s** = *A_i_β_i_* for *β_i_*, the correlation coefficient, the cosine similarity or the explained variance. In this paper we will focus on the least square solution to **s** = *A_i_β_i_*. To solve for *β_i_* we denote:

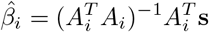

Since *A_i_* and **s** are vectors, we can rewrite 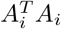 as the dot product of *A_i_* with itself and 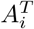 as the dot product between *A_i_* and **s**. Hence we denote:

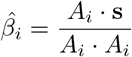

By doing so, the test statistic *η* for *f*(*x, y*) = (*x* · *y*)/(*x* · *x*) can be computed in the following way:

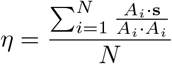

In this case *η* can be interpreted as the average fit of the data to the rank order shape. If we assume, for example, *k* = 3 groups with means [*μ_a_, μ_b_, μ_c_*] and an average rank order shape of **s** = [3, 1, 2], *η* can be interpreted as the average fit of each observation *A_i_* to that shape, or in other words, how well the data explains sorting the means, such that *μ_b_ < μ_c_ < μ_a_*. Note that we cannot state how much different each mean is from each other, but how much the data supports this rank order. As previously mentioned, *f* can be tailored to the specific needs of the respective research question. To the authors opinion however, choosing *f*(*x, y*) = (*x* · *y*)/(*x* · *x*), can be applied to a variety of research questions, and gives rise to a straightforward interpretation. Nevertheless, this specific choice for *f* is limited by the fact that if one observation *A_i_* is zero for all groups, then *f* cannot be computed and thus *η* would be invalid. Hence, *f* needs to be chosen such that it is defined for each observation and each permutation.

It needs to be pointed out that some of the proposed choices for *f* are magnitude free (i.e. correlation coefficient, cosine similarity and explained variance), whereas other choices, such as *f*(*x, y*) = (*x* · *y*)/*x* · *x*) are not. Magnitude free in this context means that the absolute value of *η* does not depend on the absolute values of either the data *A* or the values of **s**. However, due to the fact that the permutation procedure is applied in a similar fashion, the bias resulting from the absolute values of *A* (and **s**) affects the estimation of *η* and the permutation distribution equally and thus does not affect the hypothesis test itself.

So far the general idea behind obtaining the test statistic, as well as multiple possible choices for *f* have been discussed. The actual hypothesis test has been neglected so far. Generally speaking the aros test is a test on the null hypothesis (*H*_0_). As previously mentioned, this test is performed by Monte Carlo sampling from the permutation distribution under *H*_0_ and comparing the initially obtained *η* to this distribution. In a first step the number of permutations *I* is defined. *I* should be a relatively high number(e.g. *I* = 10000). Since for each permutation step an average over *N* observations needs to be computed, the maximum number of all possible permutation steps for *k* groups (and thus *k*! possible permutations per observation) can be computed as *I* = *k*!^*N*^/*N*. Where computationally feasible, the computation of the exact permutation distribution *D* under *H*_0_ (all possible combinations) should be preferred. Otherwise *I* samples (ideally, but not necessarily, without replacement [1]) need to be obtained by means of Monte Carlo sampling. During each iteration *i* ∈ [1, …, *I*], the data *A* is permuted across groups for each observation separately. Then, the test statistic is computed and added to *D*. After performing *I* iterations, *D* contains *I* values for the test statistic under *H*_0_, to which the initially obtained *η* is compared. The *p*-value is obtained by computing the fraction of all values in *D* that exceed *η*. Figure 1, depicts the aros test procedure.

**Figure 1:**
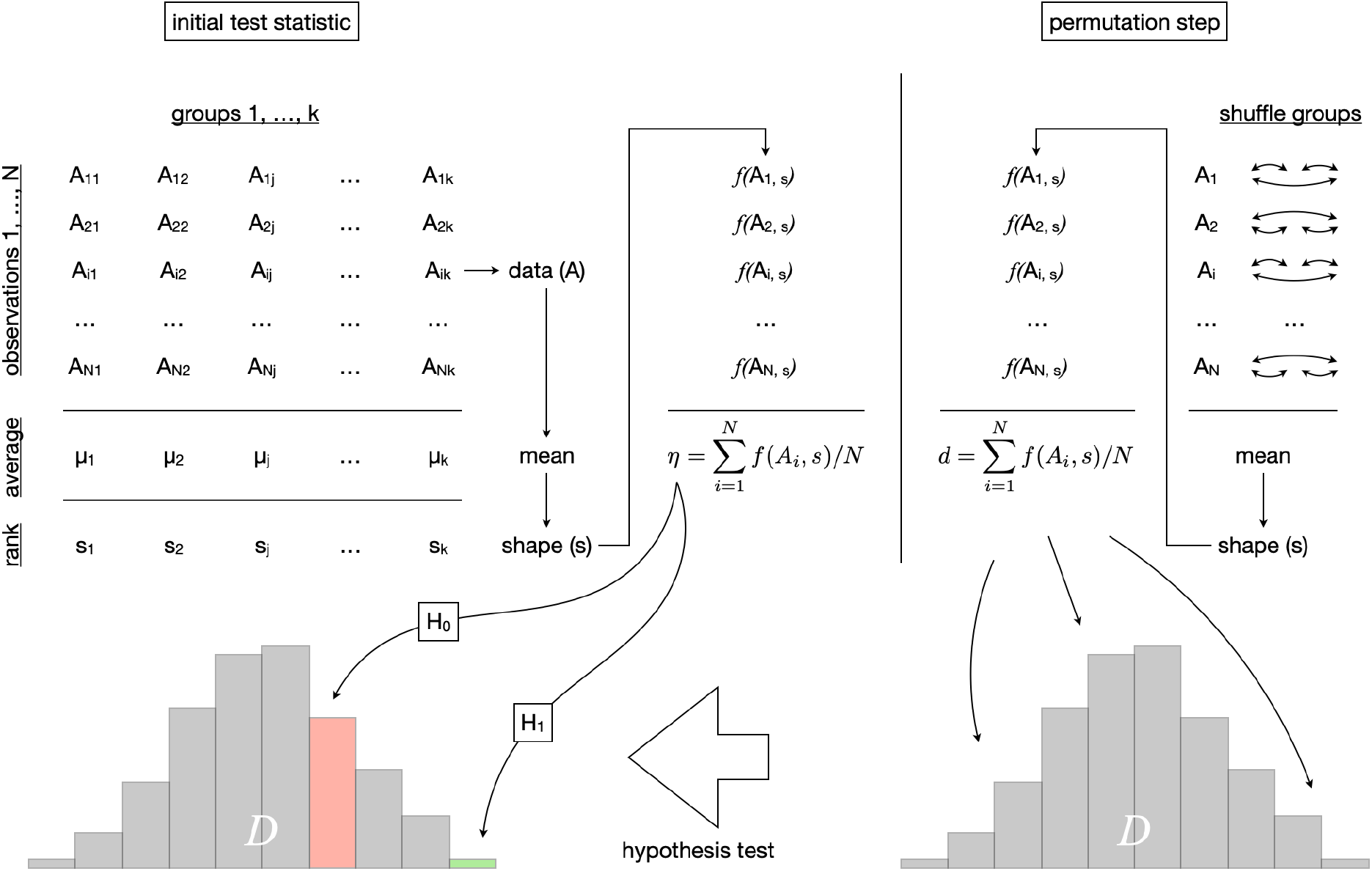
Procedure of the aros test. First, the initial test statistic *η* is obtained by averaging the data over groups and transforming the result into rank values forming the shape **s** (upper left). Afterwards, the data *A* is related to **s** using the function *f* (upper middle). This procedure is repeated *I* times to form the permutation distribution *D* (lower right), except that the data is shuffled over the data columns for each observation separately (upper right). Lastly, *η* is compared to *D* (lower left). This is achieved by computing the *p*-value as the fraction of the values in *D* that exceed *η* and compare it to the critical *α*-value (e.g. 0.05).

During the description of the test procedure, *H*_0_ has been mentioned, but was only implicitly defined. Derived from the general assumption of permutation tests, it is important that the data is exchangeable under *H*_0_. This means that the joint probability distribution (under *H*_0_) remains the same, irrespective of the order of the single values [4]. Less formally speaking, this means that if there was no difference between multiple groups of values, the values in each group could be interchanged, because they can be assumed to stem from the same distribution. The exchangeability assumption is tightly linked to *H*_0_ itself. If *η* does not depend on the grouping of the data (*H*_0_), then *η* would be expected given D, hence:

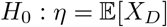

Under *H*_0_, **s** is derived from *A*, where each value in 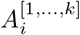 could be exchanged. Hence, **s** is the result of randomly arranged group labels and its respective average relationship with *A* - expressed by 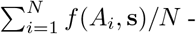 - could not be predicted by any single *A_i_* or **s**. Thus, each value within *D* could be seen as the average of independent random values. In the “Derivations” section it will be shown that *D* can be approximated by a normal distribution:

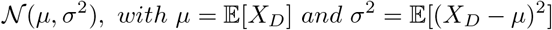

As described above, the FA rate is controlled and the hypothesis test performed by estimating the exact *p*-value 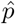 and comparing it to the critical *α*-value [1]:

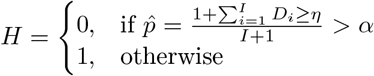

Since the *p*-value can be computed directly from *D* as the fraction of values in *D* exceeding *η* and comparing it to the critical *α*-level, the false alarm rate is controlled sufficiently.

Note that *D* can be approximated by a normal distribution (see “Derivations” for additional details) and thus *D* could be standardized: *Z* = (*D* – *μ_D_*)/*σ_D_* and *η* expressed in terms of the number of standard deviations it is different from the mean of the permutation distribution (*z*-scoring).

### A special case of aros tests

While the standard aros procedure is performed using a shape derived from the data average (data driven), a special case arises if the shape can be derived from an external source. If possible, **s** can be obtained by source *independent* of the data (e.g. previous research or derived from the hypothesis). Hence, *H*_0_ changes from asking whether a specific arrangement of the data produces a specific rank order of averages as likely as any other rank order obtained from shuffling the data, to whether under the exchangeability assumption the original arrangement of the data explains a *specific* shape better than any other arrangement of the data. Then, *η* would be computed based on this *specific* shape. Hence, the shape would not be derived from the average rank order of the data, which biases the estimation of the permutation distribution. By definition, the independence of **s** is strictly necessary. In order to avoid “double dipping” **s** cannot be derived from the mean and used as *specific* shape of interest. It must be derived from some source other than the data, in order to not violating the independence assumption of **s**. However, if **s** can be derived from some external source, not violating the independence assumption, then statistical power for this *specific* shape can greatly be increased (see “Comparison of test power”). Furthermore, it is not strictly necessary anymore to require **s** to be a unique set of equally spaced values, as the probability for any **s** to occur does not need to be uniformly distributed anymore (because there is only one shape). Technically this strips the aros test of its auto-regressive nature. Nevertheless, the general principle would remain similar enough to view this as a special case of the aros test. Please see “Discussion” for a suggested application, where the standard and special case are combined.

## 3 Derivations

The derivation is almost immediate given the statement of the central limit theorem.

### Theorem 1

(Central limit theorem[7]). *Let X*_1_, *X*_2_, … *be i.i.d. with E*[*X_i_*] = *μ*, *var*(*X_i_*) = *σ*^2^ ∈ (0, ∞). Let 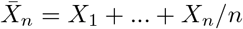. Then, as 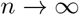,

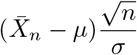

*converges to the standard normal distribution* 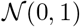

From this theorem and the definition of *η*:

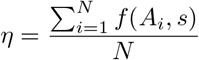

we can directly conclude:

### Corollary 1.

*Assuming that f*(*A_i_, s*) *forms a probability distribution with mean μ and variance σ*^2^, *then, for N* → ∞, *the distribution of η converges to a normal distribution* 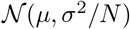

Given this result one can conclude the distribution of *η* statistics can be approximated by a normal distribution.

Now that the distribution of *η* statistics follows a known probability distribution, we can conclude that the probability distribution *f* (*X_η_* |*D*), that is the distribution of the test statistic given the permutation, is well defined. Finally, we only need to prove that in general the permutation test controls the FA rate [4]. We only need to show that the *α* level of the permutation distribution is the same as the real *α*, i.e. *P*(*reject H*_0_|*H*_0_).

### Theorem 2.

*in a permutation test, the probability α* = *P*(*reject H*_0_|*D*, *H*_0_) *is equal to P*(*reject *H*_0_|*H*_0_*)

*Proof*. We rewrite *P*(*reject H*_0_|*H*_0_) to be conditioned on *D*, as

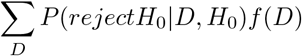

Then, by definition of the *α* level of the permutation distribution,

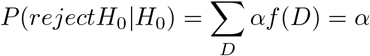

So we can conclude that, in particular, the false alarm rate of the *η* statistic permutation distribution, *f*(*X_η_*|*D*), is controlled.

## 4 Simulations

In order to demonstrate the validity and applicability of the proposed test procedure, we conducted a variety of simulations. It will be demonstrated that the type *I* error rate (FA rate) can sufficiently be controlled given the proposed procedure. Furthermore, it will be demonstrated that the permutation distribution under *H*_0_ indeed converges to a normal distribution for multiple different underlying data distributions. Additionally, one simulation will construct a specific case, where the aros test outperforms pairwise t-test and a one-way ANOVA. Lastly, it will be simulated how under specific conditions the statistical power of the aros test compares to pairwise t-tests and a one-way ANOVA. For every set of example data A, we generated 50 observation for three groups. The *α*-level was set to *α =* 0.05 and *f* was defined as *f*(*x, y*) = (*x* ·*y*)(*x* ·*x*) (see “The aros test explained”). All example simulations, as well as an implementation of the aros test are provided via https://gitlab.com/TommyClausner/aros-test.

### Estimating the type I error (FA rate)

The FA rate was estimated by 10000 simulations on uniformly distributed random data. A new data set was created for each simulation. For each test, 10000 permutations were performed to estimate the permutation distribution under *H*_0_. If the null hypothesis was rejected, the result of this simulation was set to 1 and to 0 otherwise. To obtain the final result, all individual test results were averaged, yielding a FA rate of 0.0503, which can be considered sufficiently close to the target of 0.05, as the deviation is only 0.6%. Figure 2 depicts the cumulative average of the result vector. After around 1000 simulations, the FA rate converged to 0.05, with only minor fluctuations for the other 9000 simulations.

**Figure 2:**
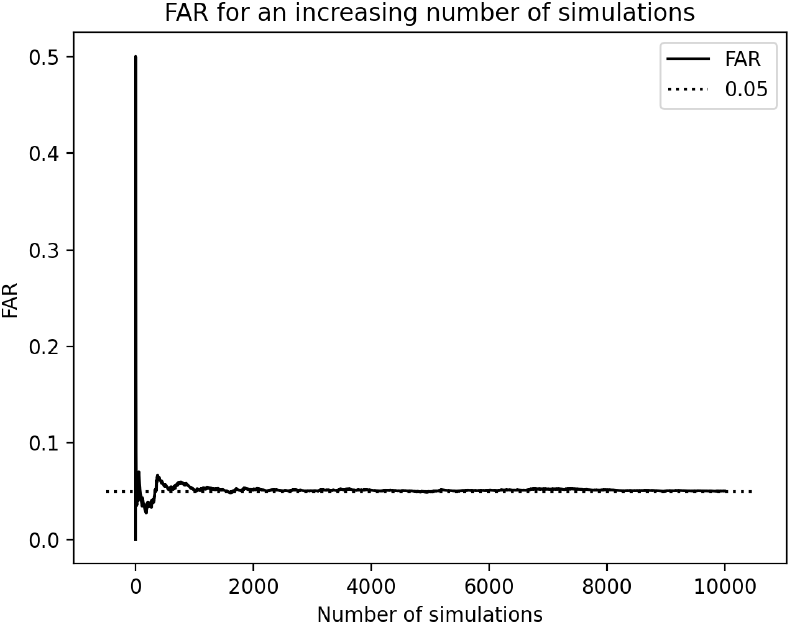
False alarm rate under *H*_0_. Each value of the continuous line represents the average of the result vector up to that respective simulation, whereas the dotted line represents the target value of 0.05 for the expected FA rate. Each value within the results vector could either be 0 (*H*_0_ was accepted) or 1 (*H*_0_ was falsely rejected).

### Demonstrating independence of sample data distributions

A crucial step towards verifying the validity of the aros test is to demonstrate that the estimated permutation distribution under *H*_0_ (the distribution *D*), indeed results in normal distribution for a variety of data distributions. As previously mentioned, *A_i_* (each paired sample) and **s** (the respective rank order shape obtained from the average over *N* observations in *A*), can be seen as independent random samples given *H*_0_. Therefore, the average of all *f* (*A_i_*, **s**) under *H*_0_ represents an average of independent random values, which according to the central limit theorem, distributes normal.

For the purpose of demonstration, four different data distributions were used to estimate the permutation distribution under *H*_0_. Random samples from the following distributions (with stated parameters), were chosen: normal distributed data with 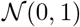, uniform distributed data with 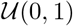, binomial distributed data with 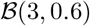 and Poisson distributed data with 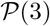. For each simulation 20000 permutations were used to estimate the permutation distribution under *H*_0_. The resulting distribution was standardized by subtracting the data to its mean and dividing it by the standard deviation. Afterwards, the data was transformed into histogram data with the number of bins determined by Sturges [8] or Freedman-Diaconis [9] rule. The approach yielding the higher number of bins was used, as is the standard for the `histogram(a, bins=’auto’)` function provided by numpy [10]. Furthermore, the count value for each bin was divided by all counts to obtain the probability for each point of the distribution. As Figure 3 clearly shows, the resulting estimated permutation distribution approximates the same normal distribution irrespective of the underlying data distribution.

**Figure 3:**
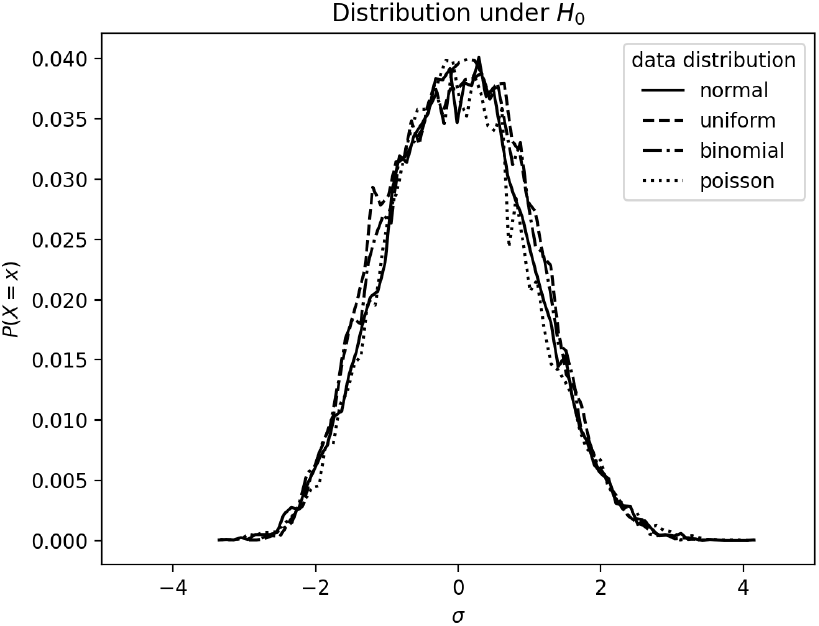
Probability distribution under *H*_0_ for a variety of input data distributions. All distributions were standardized and transformed according to *Z* = (*D* – *μ_D_*)/*σ_D_*. Afterwards the data was transformed into histogram data, where the count per bin was divided by the sum of all counts to obtain probability values for each point of the histogram. Histogram data was depicted as lines, rather than bars, for better readability.

### Simulated example

In order to compare the aros test to paired sample t-tests and one-way ANOVA, we constructed a data set in the following way: A vector of *N* = 50 random standard normal distributed values was created, to which an offset of [0.2, 0, 0.1] was added in order to simulate the difference between the group averages. Furthermore, uniform random noise was added to each group, drawn from a uniform distribution with parameters 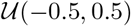. A box-plot of the simulated data for each group can be found in Figure 4. The authors point out that the same random seed as for all other simulations was used and no particular choice towards tuning the result in a desirable way was made. However, the authors are aware of the fact that changing the random seed might indeed affect the clarity of the result. However, the aim of this particular simulation was to demonstrate that there exist data sets for which the aros test is particularly well suited. Furthermore, additional 1000 simulations were conducted, where multiple sets of data using the same parameters as in this simulation were constructed and the respective test power was compared (see “Comparison of test power”).

**Figure 4:**
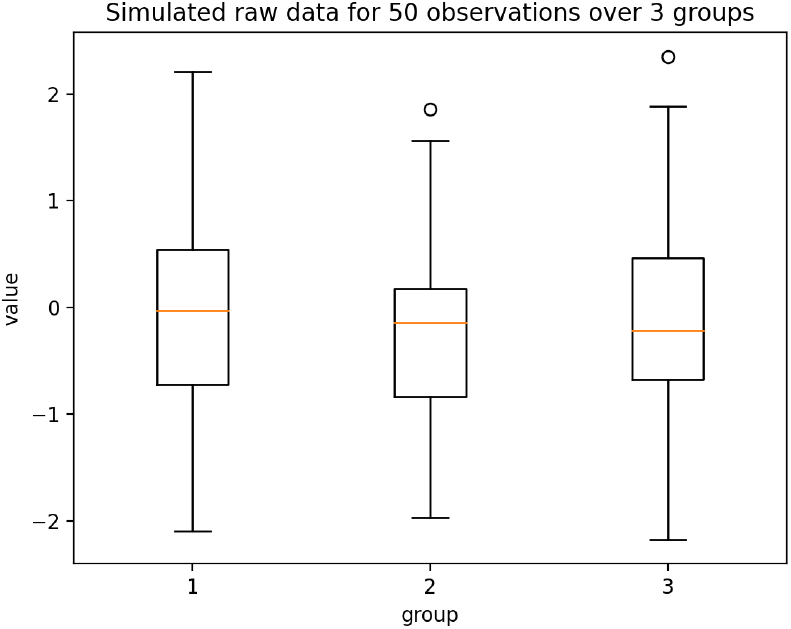
Simulated raw data. For each of the *k* = 3 groups, *N* = 50 observations were generated in the following way: *N* random samples were drawn from a normal distributions with parameters 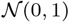. To simulate the difference in means, the same data was used three times and shifted such that *μ*_1_ = 0.2; *μ*_2_ = 0; *μ*_3_ = 0.1. Additionally uniform random noise was added separately to each group, drawn from a distribution with parameters 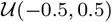.

In a first step the aros test was performed, followed by pairwise paired sample t-tests between the groups (1 vs. 2; 1 vs. 3; 2 vs. 3) and a one-way ANOVA over all three groups. The *p*-value for each test was obtained and in case of the aros test, the rank order shape as well. Due to how the data was simulated, it is expected for the three averages to be related such that *μ*_2_ < *μ*_3_ < *μ*_1_. Respective *p*-values for each test can be found in Table 1. At a critical *α*-level of *α* = 0.05, only the aros test and the t-test comparing groups 1 and 2 yielded a significant result. All other *p*-values were greater than the specified critical α-value. Additionally, from the aros test the rank order shape **s** = [3, 1, 2] could be obtained. Thus, the respective means can be ordered as *μ*_2_ < *μ*_3_ < *μ*_1_. This rank order could not have been obtained (given the data) using any other test.

**Table 1:** *p*-value of different statistical tests based on the data depicted in Figure 4

| test | $p$-value |
| --- | --- |
| aros test | 0.001 |
| t-test 1 vs. 2 | 0.003 |
| t-test 1 vs. 3 | 0.065 |
| t-test 2 vs. 3 | 0.203 |
| one-way ANOVA | 0.618 |

### Comparison of test power

In order to approximate the relative test power given the scenario explained in “Simulated example”, the procedure was repeated 1000 times (with similarly constructed, but newly generated data), where we recorded the number of correctly rejected *H*_0_ for each test scenario. The following scenarios were included: aros test (aros), aros test with known (pre-determined) shape (*aros^ks^*), pairwise t-tests with uncorrected *p*-values (*t^UC^*), pairwise t-tests with Bonferroni (*t^B^*) or Bonferroni-Holm (*t^BH^*) correction and a one-way ANOVA (*ANOVA*) for simultaneous comparison of all groups. During each of the 1000 simulations, it was recorded whether *H*_0_ was correctly rejected or not. For the t-test scenarios, *H*_0_ was counted as correctly rejected, if *p*-values of all three tests (corrected or not), were lower than or equal to the critical *α*-level, which was set to *α* = 0.05. For all other test tests, *H*_0_ was counted as rejected, if the respective *p*-value was lower than or equal to *α* and the predicted shape corresponded to the presumed shape from which the data was constructed. This last condition was of cause omitted from the ANOVA scenario. In a last step, the cumulative average of each condition was divided by the overall average of the uncorrected t-test scenario, which acted as a baseline. Figure 5 depicts the cumulative, relative average for each condition. Thereby a change of the *y*-axis, e.g. by a factor of two, would indicate that the relative test power of the respective test was twice as high as the estimated test power of the uncorrected t-test after 1000 simulations.

**Figure 5:**
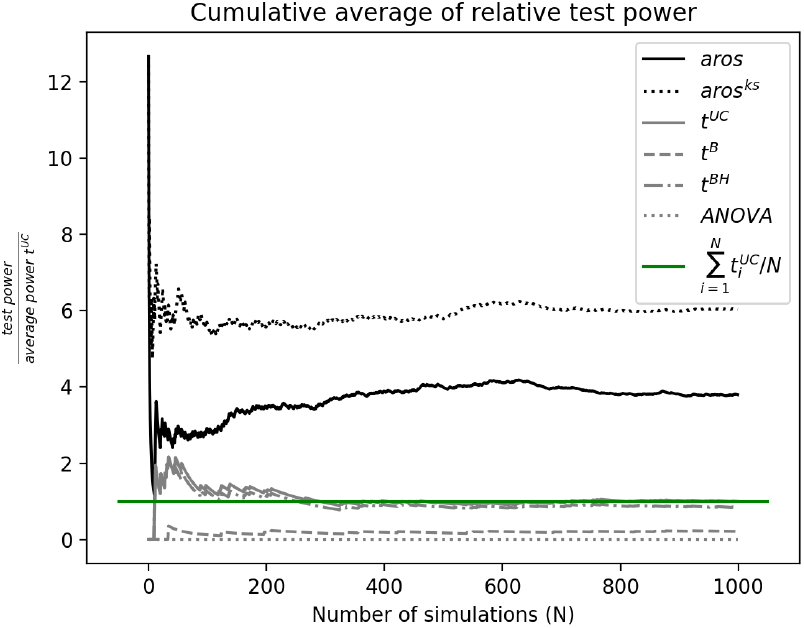
Cumulative average of relative test power. For each condition, the number of correctly rejected *H*_0_ was counted and plotted cumulatively, relative to the number of simulations. This was done for *k* = 3 groups. Additionally, each value was divided by the final average of correctly rejected *H*_0_ obtained from the uncorrected t-test scenario (*t^UC^*). For each of the t-test scenarios, *H*_0_ was counted as successfully rejected if all three of the pairwise comparison yielded a *p*-value lower than or equal to the critical *α*. This procedure was performed for the regular aros test (*aros*), a version of the aros test, where the shape was set to be known (*aros^ks^*), pairwise t-test without correction for MCP (*t^UC^*) and with correction for MCP using the Bonferroni (*t^B^*) or the Bonferroni-Holm method (*t^BH^*). Additionally a one way ANOVA for the simultaneous comparison of all three groups was performed (*ANOVA*). The green line indicates the final average of t^UC^ after 1000 iterations.

None of the traditional methods was able to capture the respective difference between the three averages as good as the aros test. Note that the result for the aros test with known shape (*aros^ks^*) needs to be taken with a grain of salt. Technically **s** was not derived from an external source and must be considered “double dipping” in this scenario. Nevertheless, it was included to demonstrate the potential increase in statistical power for that case.

## 5 Discussion

We propose a paired samples, non-parametric statistical test procedure on the basis of a permutation test, which aims to circumvent the MCP by combining the relationship between multiple group averages into a single statistical value. Via an autoregressive procedure, paired sample data for multiple groups is related to the rank order shape of the group averages. As such, the aros test acts as a test for interval scaled data, where the test result needs to be interpreted on the basis of ordinal scaled ranks. It was demonstrated that the FA rate is controlled sufficiently by a permutation procedure, where the estimated permutation distribution *D* under *H*_0_ can be described by a normal distribution with parameters 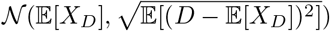. A set of simulations has been conducted to verify the procedure. First, the FA rate was estimated over 10000 simulations, which yielded an estimated value of 0.0503. This value can be considered sufficiently close to the target value of 0.05. In order to save computation time, each simulation was based on 10000 permutations and *k* = 3 groups.

Since it could be demonstrated that each *f*(*A_i_*, **s**) yields an independent random value, the authors do not assume the FA rate to change on the basis of the number of groups, nor by increasing the number of permutations per simulation. Furthermore, simulations on the basis of multiple data distributions showed that the estimated permutation distribution under *H*_0_ distributes normal, as predicted. It needs to be pointed out that no particular reason for the choices of the data distributions can be brought forward. Instead - to the authors experience - the most common distributions have been chosen. Again, since each value in *D* stems from an average of independent random values, it is not expected that different choices for the data distributions would affect the normality of the estimated permutation distribution under *H*__0__. In a third simulation, the aros test was compared to the probably most common statistical tests: the t-test and ANOVA. Since the purpose of this paper was to demonstrate that conducting an aros test in some circumstances can be beneficial where t-tests or ANOVAs fail, the data was deliberately constructed such that the result would favor the aros test. However, no particular manipulation was applied to the data. Instead the data was constructed based on normal distributed data (required by t-test and ANOVA), where the difference in means was marginal compared to the noise that was added. This particular case (low signal-to-noise ratio) is one example where the aros test potentially outperforms classical tests. Nevertheless, the aros test has not been applied to a real data set in the current publication, which needs to be addressed in the future. However, one of the authors can confirm that the aros test has been applied successfully to a set of (yet unpublished) neuro-scientific data. In a last simulation, the relative power between the aros test and traditional tests has been assessed. For 1000 simulations, the true positive rate was compared between the tests. Thereby, the true positive rate of uncorrected t-tests served as baseline and the result was computed relative to that average. Since the statistical power for tests on the null hypothesis is generally hard to assess and is strongly dependent on the data, the relative power to well established statistical tests might provide insight into how the aros test performs under specific circumstances. On average, the aros test performed more than three times as good as a uncorrected t-test, given the respective data. However, it needs to be pointed out that positive results for the pairwise t-tests were only counted if all three tests yielded a significant result. This procedure might be overly conservative in many circumstances. However, in this specific case, the distribution of the group averages was known beforehand and the goal was to determine the specific order in which those averages could be sorted. For such a statement to be made using t-tests, indeed three significant results would be required. Since not a single test using the one-way ANOVA yielded a significant result, the authors presume that it would not be suited for the respective data set. Relating within and between subject variance might have failed due to the large overall variability in the data that was much greater within than between the groups. Lastly, the aros test with pre-defined, known shape performed almost six times better then the baseline method. Since the present example would clearly be a case of “double dipping”, this result should not be overstated. However, in a real world scenario, if the a specific shape would be expected (e.g. due to how the hypothesis was constructed or by previous research), this method potentially increases statistical power significantly.

In general, the biggest advantage of the aros test is its capability of allowing for a qualitative conclusion about the relationship between more than two group averages without prior knowledge of the underlying data distributions. Thereby, the test is relatively robust against noise. Depending on the choice for *f*, a significant result can be interpreted in multiple ways. Irrespective of *f* however, rejecting *H*_0_ means that the paired observations [1, …, *N*], are not fully independent and that the shape **s** can be predicted by *A* with a probability *P*(**s**) > 1/*k*!. This means that the labels of groups [1, …, *k*] are not exchangeable since the distributions for *D*_[1, …, *k*]_ are different. In other words, grouping the data into [1, …, *k*] groups with labels [1, …, *k*] is in fact meaningful and can be justified by the data. The respective choice for *f* enriches this finding with some additional information. If *f* was set to e.g. *f*(*x, y*) = (*x* · *y*)(*x* · *x*), *η* indicates how many units of change in the data explain a single unit of change in the shape vector **s**.

As mentioned before, the aros test is meant as an alternative for a variety of scenarios where more than two groups need to be compared, but classical statistical tests fail, either by violation of assumptions (e.g. violation of normality of the underlying data distributions) or if the MCP reduces statistical power unsatisfactorily. We could further demonstrate that the aros test potentially outperforms classical tests in low signal-to-noise ratio scenarios. Another positive aspect is the additional information that can be obtained by the rank order shape. An ANOVA, applied to multiple groups, can inform the user about whether there is any difference across the tested groups, whereas the aros test additionally provides information about the qualitative relationship between the group averages. However, the relatively high statistical power to obtain an ordinal result from interval scaled data, circumventing the MCP, comes at a cost.

First and foremost the aros test does not provide any information about the quantitative difference between a set of means and can furthermore not be interpreted in such a way. Thus, it is impossible to obtain any effect size for the differences in means and hence should only be applied with this knowledge in mind. In some way the absolute difference between group averages is sacrificed in favor of an increased power for relative differences between the groups. Second, the result can only be interpreted in its entirety. This means no sub-comparisons between the relationship of means can be made and only the entire rank order profile as such can be interpreted. Note that this is similar to the result of a cluster based permutation test [4]. This leads directly to a third caveat, that is the number of groups that can be meaningfully compared. While the interpretation of the rank order shape for three groups in most cases might be straight forward, it might not be for a large number of groups. In general, if a very large number of groups is compared and at least one dimension of the data is correlated, it might be advisable to choose a cluster permutation test [4]. Since it is possible to compute the aros test for an arbitrary large number of groups, the authors can only provide a rule of thumb, based on their experience in the fields of experimental Psychology and Neuroscience. The authors believe that the number of groups *k* to be compared, should be kept in a range where 3 ≤ *k* ≤ 7. Values for *k* higher than that might be extremely difficult to interpret. Moreover, if *H*_0_ was rejected, the obtained shape can only be interpreted if the group averages are indeed unique. This means that in order for the aros test to be interpretable, it needs to be ensured beforehand that each group average is numerically unique.

Lastly, the authors would like to point towards a variation of the aros test, which can be considered similarly valid as the standard procedure. However, if not applied carefully, this variation quickly leads to a circular analysis (“double dipping”). In some scenarios, the expected shape that is explained by the data might already be known. This can be the case if previous experiments allow for a justified prediction or in other cases the experimental hypothesis pre-determines the expected shape already. However, it needs to be emphasized that irrespective of the origin of the shape, it needs to be *independent* to not bias the estimation of the permutation distribution under *H*_0_. If such an independent shape could be derived from some external source other than the data, then this shape could be used to obtain *η* and computing *D*. Hence, one could test whether a *specific* rank order can be justified by the data with a probability higher than 1/*k*!. Additionally, this approach can be combined with the classical procedure: If two independent data sets with *k* groups exist, then the classical approach could be applied to one of the data sets and - in case *H*_0_ was rejected - the obtained shape can be tested against the second data set.

## 6 Conclusion

As demonstrated, the auto-regressive rank order similarity (aros) test can be considered an alternative test to circumvent the MCP for a small number of groups in a paired sample statistical test setting. While the ability to determine the magnitude of differences between group averages is lost, additional statistical power is gained to test the relationship between the raw data and the rank order of the group averages. Since the aros test is based on a permutation procedure to estimate the permutation distribution under *H*_0_, no assumptions about the distribution of the data are required other than exchangeability under *H*_0_. Furthermore, it has been demonstrated that the aros test controls the FA rate sufficiently. Since the aros test relates the magnitude of the group averages, without comparing them directly, it is exceptionally well suited for test scenarios, where the signal-to-noise ratio is low and the rank order of the means is of higher interest than the actual effect size.

## Acknowledgments

The authors would like to express their gratitude towards Mathilde Bonnefond, Jérémie Mattout and René Scheeringa for fruitful discussions and general support.

## Footnotes

1 There is a large variety of strategies, which are not discussed here.

